# Ameliorative effect of a quinazoline-based bromodomain inhibitor, CN210, on experimentally-induced Crohn’s disease-like murine ileitis via inhibition of inflammatory cytokine expression

**DOI:** 10.1101/2020.01.24.917948

**Authors:** Takehisa Noguchi, Kyosuke Hidaka, Satsuki Kobayashi, Kenjiro Matsumoto, Makoto Yoshioka, Shyh-Ming Yang, David J. Maloney, Shinichi Kato

**Affiliations:** Division of Pathological Sciences, Department of Pharmacology and Experimental Therapeutics, Kyoto Pharmaceutical University, Kyoto, Japan; ConverGene LLC, 3093 Beverly Lane, Unit C, Cambridge, MD 21613, USA; National Center for Advancing Translational Sciences, National Institutes of Health, 9800 Medical Center Drive, Rockville, MD 20850, USA

## Abstract

Inhibitors of bromodomain and extra-terminal motif proteins (“BET inhibitors”) are emerging epigenetic therapeutics that exert their effects by suppressing the expression of genes that drive cancer and inflammation. The present study examined the anti-inflammatory effects of a quinazoline-based BET inhibitor, CN210, which also inhibits bromodomains of two paralogous histone acetyltransferases (HATs), such as cAMP-responsive element binding protein-binding protein (CBP) and p300. To assess its effectiveness against inflammatory bowel disease (IBD), CN210 was tested in an experimentally-induced murine Crohn’s disease (CD)-like ileitis model. Ileitis was induced in mice by subcutaneous administration of indomethacin and CN210 was given orally 30 min before and 24 h after the indomethacin administration. Whilst indomethacin produced severe ileitis accompanied by an increase in ileal mucosal myeloperoxidase (MPO) activity, administration of CN210 reduced the severity of ileitis in a dose-dependent manner and suppressed the MPO activity. Similarly, upregulation of inflammatory cytokines was observed in ileitis mucosa after indomethacin injection but this response was significantly attenuated by CN210 administration. To further characterize the effects of CN210 on inflammatory pathways, monocyte/macrophage-like cell line RAW264 treated with lipopolysaccharide (LPS) was exposed to CN210. CN210 significantly attenuated the expression of inflammatory cytokines and reversed the activation of NF-κB and MAP kinases induced by LPS. These findings suggest that CN210 ameliorates indomethacin-induced ileitis due at least in part to the inhibition of inflammatory cytokine expression *via* attenuation of NF-κB and MAP kinase pathways, thus representing a new mode of therapy for CD.

## Introduction

Bromodomain and extra-terminal motif (BET) proteins, including BRD2, BRD3, BRD4 and BRDT, are epigenetic ‘reader’ proteins that bind to acetylated lysine residues on proteins, including histones, by means of their bromodomains [1, 2]. Of the four BET proteins, BRD4 associates with general transcription cofactor complex ‘Mediator’ and is a key structural component of extensive transcription factor complexes formed at the genomic regions termed Super-Enhancers (SEs) [2–4]. Small-molecule inhibitors of BRD4 was shown to suppress the expression of MYC oncogene and the genes that encode pro-inflammatory cytokines, such as IL-6, IFN-β, IL-1β, IL-12α, CXCL9 and CCL12 in mice [5–7], and showed efficacy in a wide range of cancer models including pancreatic, breast, ovarian, and colon cancers [8–12]. Accordingly, small molecule inhibitors of BET proteins (“BET inhibitors”) are now considered as promising drug candidates for both cancer and inflammatory diseases [13, 14].

Whilst several BET inhibitors currently are evaluated clinically [15], early results showed mixed results with undesirable side effects including thrombocytopenia [16]. Therefore, further development of alternative BET inhibitors with distinct structures and better safety and efficacy profiles remains of high interest. We recently developed a novel quinazoline-based BET inhibitor CN210 (Fig 1A) with a therapeutically suitable target affinity (BRD4 BROMOscan^®^ K_d_: 70 nM), cellular potency (IC_50_ of 0.94 µM for the viability assay in leukaemia cell line, MV4-11) and a pharmacokinetic profile (t_1/2_ of 1.7 h; C_max_ of 2,163 ng/mL; AUC_0–∞_ of of 8,927 ng·h/mL; and oral bioavailability of 75% following a 10 mg/kg oral administration in mice) [17]. CN210 is highly effective in the inhibition of tumour growth in Kasumi-1 xenograft mouse model and in the improvement of arthritis in collagen-induced mouse arthritis model [17].

**Fig 1.**
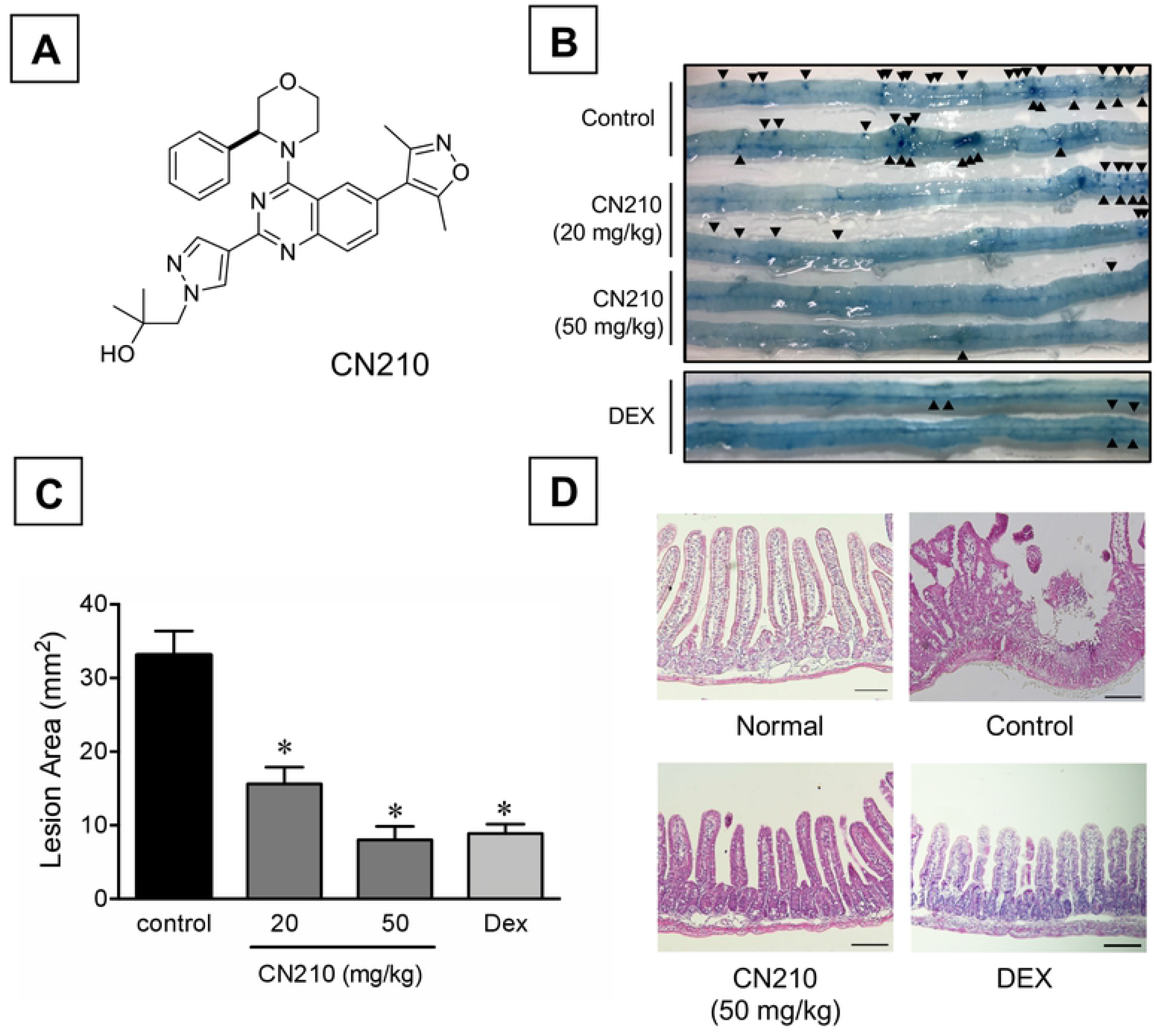
Effect of CN210 on indomethacin-induced ileitis. Animals received indomethacin (10 mg/kg) subcutaneously and sacrificed 48 h later. CN210 (20 and 50 mg/kg) and dexamethasone were administered 30 min before and 24 h after indomethacin treatment. Chemical structure of quinazoline-based BET inhibitor CN210 (A). Representative macroscopic observation of indomethacin-induced ileitis (B). Arrow heads indicate Evans blue-stained ileal lesions. Lesion area of indomethacin-induced ileitis (C). Data are shown as the mean ± SEM (n = 5–8). Statistically significant difference at \**P* < 0.05 from control (vehicle alone) analyzed by one-way ANOVA with Holm-Sidak’s test. Representative histological observation of indomethacin-induced ileitis at x100 magnification (D). Scale bar: 50 µm.

Inflammatory bowel disease (IBD), including ulcerative colitis and Crohn’s disease (CD), is characterized by chronic inflammation and damage in gastrointestinal tracts resulting from abnormal immune responses and aberrantly increased inflammatory cytokines. Whilst recent introduction of biologic agents, such as anti-TNF-α and anti-IL-12/23p40 monoclonal antibodies, has significantly improved the outcome of IBD [18–20], complete remission still is a challenge as IBD patients often develop anti-drug antibodies diminishing the long-term effectiveness of these agents. Thus, development of small-molecule drugs that complement current armamentarium to treat IBD is of high medical interest.

Recent studies showed that BET inhibitors ameliorates colitis by inhibiting the differentiation of Th17 cells, maturation of dendritic cells and suppression of inflammatory cytokine expressions due at least in part to the blocking of NF-κB and MAPK activation [21–23].

Non-steroidal anti-inflammatory drugs (NSAIDs), including indomethacin, produce damage not only in the stomach but also in small intestine in humans and experimental animals [24–26]. Because of the pathomorphological similarity to human CD in small intestine, rodent indomethacin-induced ileitis model is considered to be a useful model of human CD [27].

In the present study, we showed that, CN210 not only inhibited the bromodomains of BET proteins but also the bromodomains of cAMP-responsive element binding protein-binding protein (CBP) and p300, two paralogous histone acetyl transferases (HATs). We also showed that CN210 suppressed in indomethacin-induced human CD-like ileitis in mice with concomitant suppression of gene expression of pro-inflammatory cytokines.

## Materials and Methods

### Drugs

Indomethacin and dexamethasone were obtained from Sigma-Aldrich (St. Louis, MO, USA). CN210 was synthesised by Dr. Shyh-Ming Yang (the National Center for Advancing Translational Sciences (NCATS), National Institute of Health) [17]. Indomethacin was dissolved in physiological saline with a drop of Tween 80 (Wako, Osaka, Japan) and administered subcutaneously in a volume of 0.1 mL/10 g body weight. CN210 and dexamethasone were suspended in carboxymethylcellulose (CMC, Nacalai Tesque, Kyoto, Japan) and administered orally in a volume of 0.1 mL/10 g body weight.

### Competitive ligand binding assay

Competitive ligand binding assays against CBP and p300 proteins were performed with BROMOscan^®^ service at Eurofins DiscoverX Corporation (San Diego, CA). Briefly, T7 phage displaying bromodomains of CBP or p300 was grown in parallel in 24-well blocks in an *E. coli* host derived from the BL21 strain: *E. coli* were infected with T7 phage and incubated with shaking at 32°C until lysis (90-150 minutes). The lysates were then centrifuged (5,000 x g) and filtered (0.2 µm) to remove cell debris. To generate affinity resins for the assays, streptavidin-coated magnetic beads were treated with biotinylated small molecule or acetylated peptide ligands for 30 minutes at room temperature. The liganded beads were blocked with excess biotin and washed with SEA BLOCK Blocking Buffer (Thermo Fisher, Rockford, IL), 1 % BSA, 0.05 % Tween 20, 1 mM DTT) to remove unbound ligand and reduce non-specific phage binding. Binding reactions were assembled by combining DNA-tagged protein, liganded affinity beads, and test compounds in 1x binding buffer (17% SeaBlock, 0.33x PBS, 0.04% Tween 20, 0.02% BSA, 0.004% Sodium azide, 7.4 mM DTT). Test compounds were prepared as 1000x stocks in DMSO and subsequently diluted to ensure a final DMSO concentration of 0.1% DMSO. The assay plates were incubated at room temperature with shaking for 1 hour and the affinity beads were washed with wash buffer (1x PBS, 0.05% Tween 20). The beads were then re-suspended in elution buffer (1x PBS, 0.05% Tween 20, 2 µM non-biotinylated affinity ligand) and incubated at room temperature with shaking for 30 minutes. The DNA-tagged protein concentrations in the eluates were then measured by qPCR. K_d_ values were obtained using a 3-fold serial dilution across 11 compound concentrations ranging from 0 µM to 10 µM and by calculating a standard dose-response curve using Hill equation with a slope of −1. Curves were fitted using a non-linear least square fit with the Levenberg-Marquardt algorithm.

### Animals and ethics statement

This study was carried out in strict accordance with the recommendations found in the Guide for Care and Use of Laboratory Animals (8^th^ Edition, National Institutes of Health). The protocols were approved by the Committee on the Ethics of Animal Research of Kyoto Pharmaceutical University (Permit Number: PETH-19-004). Male C57BL/6 mice weighing 22–26 g (SLC Co., Shizuoka, Japan) were acclimated to standard laboratory conditions with 12-h light–dark cycles and a temperature of 22 ± 1 °C. Experiments were performed using five to eight un-anaesthetised mice per group.

### Induction of ileitis

After fasting for 18 h and refeeding for 1 h, the animals were administered subcutaneously 10 mg/kg of indomethacin and sacrificed 48 h later. CN210 (20 and 50 mg/kg) and dexamethasone (3 mg/kg) were administered orally 30 min before and 24 h after indomethacin injection. The small intestine was removed and fixed with 2% formalin. The fixed small intestine was opened along the anti-mesenteric attachment and examined for lesions under a dissecting microscope (S6D, Leica Microsystems, Wetzlar, Germany) (x20). The area of macroscopically visible lesions (mm^2^) was measured and summed per ileum. To visualise lesions, 0.5% of Evans blue solution was injected intravenously in a volume of 0.2 mL/animals 30 min before sacrifice. The measurement of area was performed by two investigators blinded to experimental groups. Subsequently, the ileum was immersed in 10% formalin overnight, and tissue samples were excised, embedded in paraffin, sectioned into 4-µm slices, and stained with haematoxylin/eosin. Histological ileitis was observed under a light microscope (BX-51, Olympus, Tokyo, Japan) fitted with a digital camera system (DS-Ri1, Nikon, Tokyo, Japan).

### MPO activity

Ileal tissues were rinsed with cold phosphate-buffered saline, weighed, and homogenised in 50 mM phosphate buffer pH 6.0 containing 0.5% hexadecyltrimethylammonium bromide. Total protein in lysates was estimated using a spectrophotometric assay kit (Pierce, Rockford, IL, USA), whereas myeloperoxidase activity was determined using *o*-dianisidine hydrochloride (Sigma-Aldrich), as previously described [53].

### Cytokine expression

Ileal tissues were washed with cold phosphate-buffered saline and stored in RNAlater (Ambion, Austin, TX, USA) at 4 °C until use. Total RNA was extracted using Sepasol-RNA I Super G (Nacalai Tesque) according to the manufacturer’s instructions and reverse-transcribed using PrimeScript Reverse Transcriptase (Takara Bio, Shiga, Japan). Expression of cytokines was quantified using a Thermal Cycler Dice Real Time System (Takara Bio) with SYBR Premix ExTaq II (Takara). Pre-designed primer sets for mouse TNF-α (primer set MA097070), IL-6 (primer set MA039013), IFN-γ (MA025911), IL-17 (MA157056), and TATA-binding protein (TBP) (MA050367) were obtained from the Perfect Real-Time Supporting System (Takara Bio). The mRNA expression level was standardised to that of TBP.

### Cell culture

Mouse monocyte/macrophage cell line RAW264 were obtained from RIKEN BRC (Ibaraki, Japan). The cells were grown in Dulbecco’s modified Eagle medium (DMEM: Wako) containing 10% fetal bovine serum (FBS) with 100 U/mL of penicillin G and 100 µg/mL of streptomycin at 37 °C in 5% CO_2_ with a humidified atmosphere. Sub-confluent cells in 12-well plate were exposed of lipopolysaccharide (LPS) (Sigma-Aldrich) at a concentration of 1 µg/mL and total RNA was extracted 4 h later. Subsequently reverse transcription and PCR were performed as described above. Pre-designed primer sets for mouse TNF-α (primer set MA097070), IL-6 (primer set MA039013), IL-12A (MA0287533), IL-23A (MA095159), and TATA-binding protein (TBP) (MA050367) were obtained from the Perfect Real-Time Supporting System (Takara Bio). The mRNA expression level was standardised to that of TBP. CN210 at 0.1, 1 and 10 μM and dexamethasone at 1 μM were co-treated with LPS in RAW264 cells.

### Western blotting

Nuclear proteins were isolated from RAW264 cells using Nuclear Extract Kit (Active Motif, Carlsbad, CA, USA) according to the manufacturer’s protocols. Total proteins were extracted from RAW264 using radioimmunoprecipitation assay (RIPA) buffer containing protease inhibitors (Complete Mini, Roche Diagnostics GmbH, Mannheim, Germany) and phosphatase inhibitors (PhosSTOP EASYPack, Roche Diagnostics) as described previously [54]. Nuclear and total protein samples were separated on sodium dodecyl sulphate-polyacrylamide gel (SDS-PAGE) electrophoresis and subsequently transferred to polyvinylidene difluoride (PVDF) membrane (Bio-Rad, Hercules, CA, USA). After blocking using 5% of bovine serum albumin (BSA) or skim milk for 1 h at room temperature, the specific primary antibodies for NF-κB, p65, p38, phospho-p38, ERK, phospho-ERK, JNK, phosphor-JNK (Cell Signaling, Danvers, MA,USA), lamin B1 (ProteinTech, Chicago, IL, USA), and β-actin (Gene Tex, Irvine, CA, USA) were treated with the PVDF membranes at 4 °C overnight. Subsequently, the membranes were incubated with horseradish peroxidase-conjugated secondary antibodies (Cell Signaling) for 1 h at room temperature, and protein expression signals were detected using a chemiluminescence detection kit (Perkin Elmer, Waltham, MA, USA) and visualised using a chemiluminescence imaging system (Fusion Solo S, Viber, Collegien, France). The levels of protein expression were normalised to that of lamin B1 or β-actin.

### Statistical analysis

All the data are shown as mean ± SEM. Statistical analysis was performed with one-way analysis of variance (ANOVA) followed by a Holm-Sidak’s multiple comparison test. *P* < 0.05 was considered statistically significant. The GraphPad Prism 6.0h (GraphPad Software, La Jolla, CA, USA) was used for statistical analysis.

## Results

### CN210 revealed competitive binding effects for the bromodomains of CBP and p300

We previously reported that CN210 potently inhibited bromodomains of BET proteins with K_d_ values ranging from 11 nM to 200 nM [17, 28]. In the present study, we demonstrated that CN210 also inhibited the bromodomains of two paralogous HATs, CBP and p300, with K_d_ values of 260 nM and 190 nM, respectively (Table 1) as compared to the previously reported K_d_ values of 70 nM for BRD4 [17]. It should be noted that CBP and p330 are proximal to BET proteins in a phylogenetic tree generated with three-dimensional structure-based alignments [1].

**Table 1.**
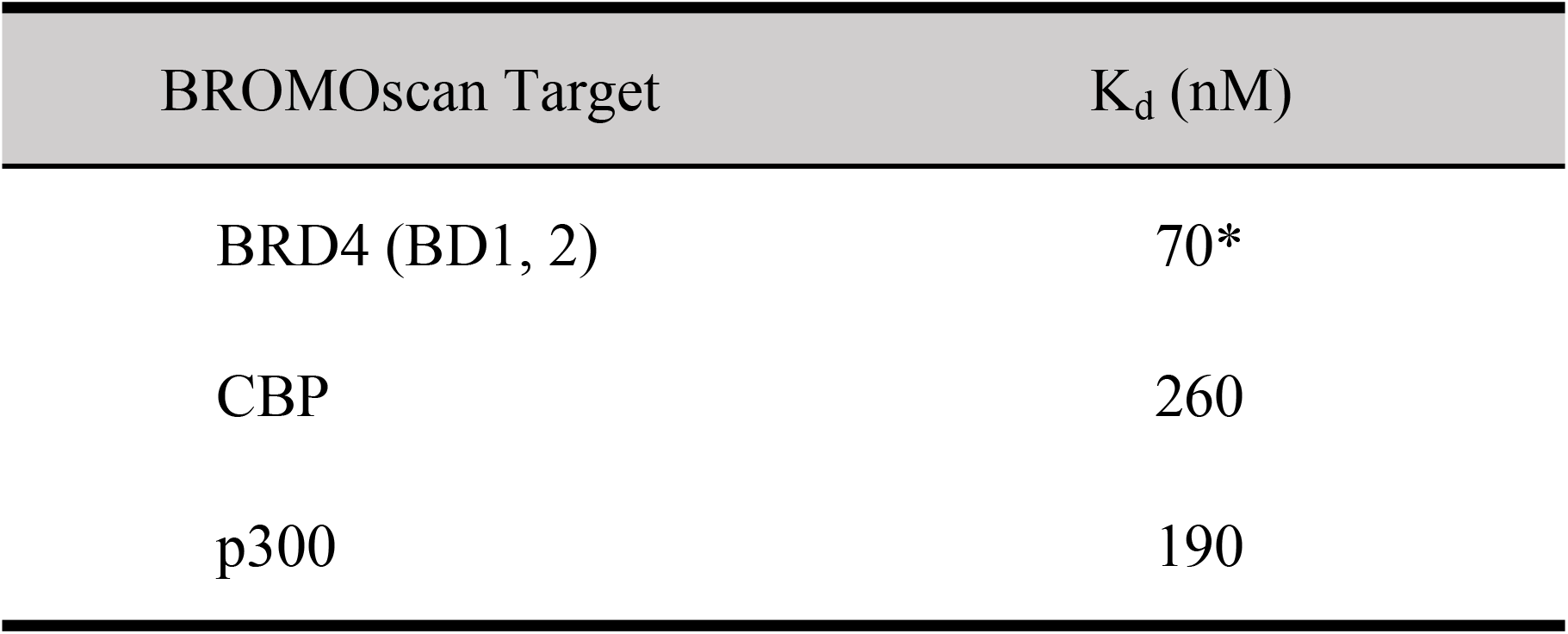
The competitive binding affinity (K_d_) for BRD4, CBP and p300.

The Competitive ligand binding assay was conducted using BROMOScan^®^ service (Eurofins DiscoverX, San Diego, CA, USA). BROMOscan was conducted against bromodomain in histone acetyltransferases, CBP and p300. *BRD4 (BD1, 2) data was from our previous study [17] and is shown for comparison.

### CN-210 reduced the severity of indomethacin-induced ileitis

Single subcutaneous administration of indomethacin produced ileitis, which is characterized by severe lesions in the small intestine, mostly in the distal jejunum to ileum (Fig 1B). Whilst CN210 (20 and 50 mg/kg) given orally 30 min before and 24 h after indomethacin administration reduced the severity of ileal lesions in a dose-dependent manner, significant effect was observed at a dose of 20 mg/kg (Figs 1B and C). Likewise, dexamethasone (3 mg/kg), a reference corticosteroid, given orally 30 min before and 24 h after indomethacin administration significantly attenuated the ileitis. Histological observation indicated that severe mucosal lesion reaching the muscle layer was found 48 h after indomethacin administration (Fig 1D). In contrast, such lesion was attenuated by the treatment with either CN210 (50 mg/kg) or dexamethasone (3 mg/kg), resulting in mucosa that appeared mostly normal.

### CN210 suppressed indomethacin-induced increase in MPO activity and cytokine expression in the ileal mucosa

To confirm the suppressive effect of CN210 on indomethacin-induced ileitis, we examined the effects of CN210 on MPO activity and cytokine expression in the ileitis mucosa. Single subcutaneous administration of indomethacin markedly increased MPO activity in the ileal mucosa (Fig 2). This increase was suppressed significantly by CN210 (50 mg/kg) and dexamethasone (3 mg/kg). Concomitant to the increase in the activity of MPO, gene expression of cytokines such as TNF-α, IL-6, IFN-γ, and IL-17 in the ileal mucosa was markedly enhanced at 48 h after indomethacin administration (Fig 3). These responses were abolished by the treatment with CN210 (50 mg/kg) or dexamethasone (3 mg/kg).

**Fig 2.**
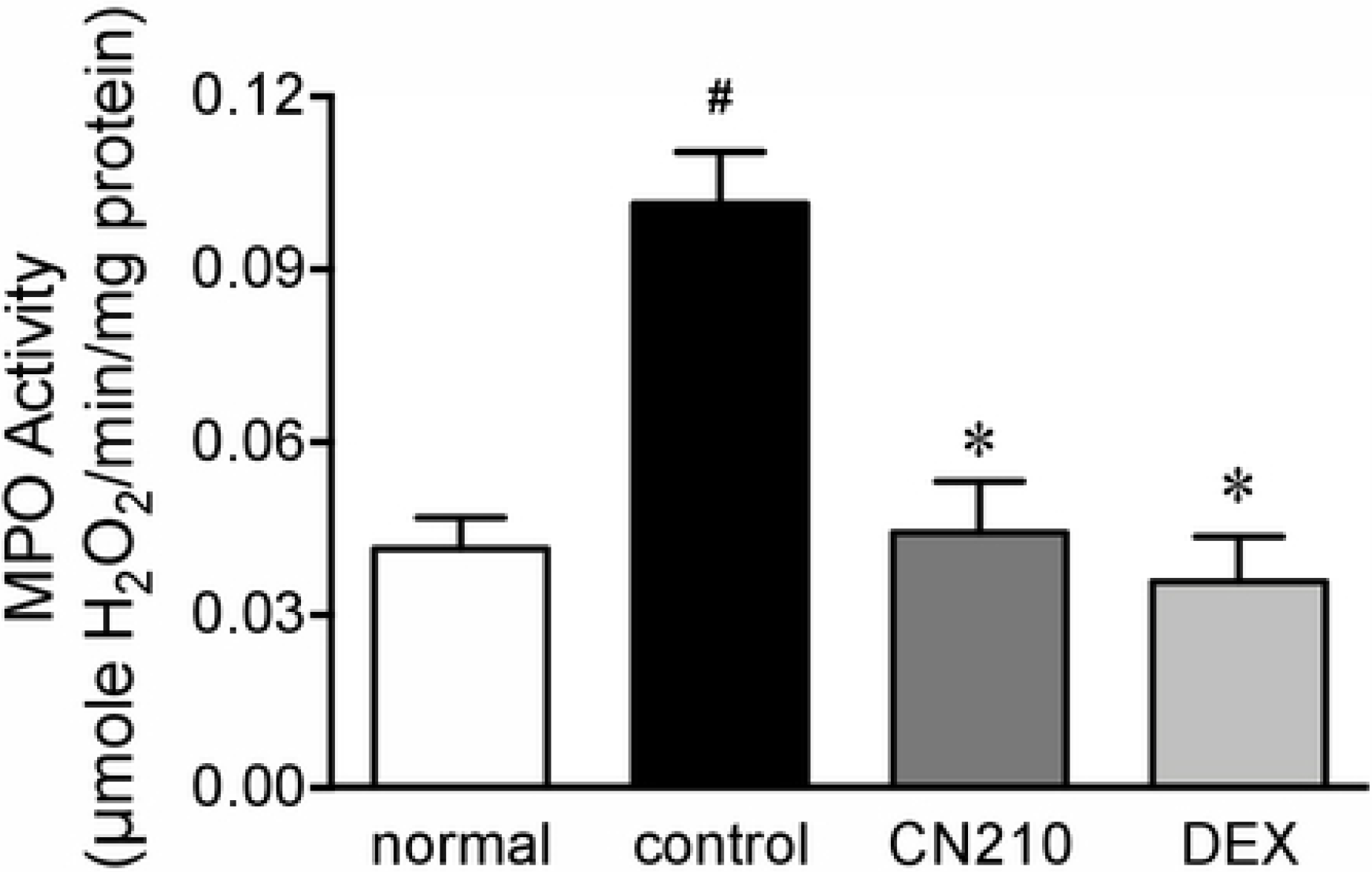
Effect of CN210 on indomethacin-induced increase in MPO activity in the ileal mucosa. Animals received indomethacin (10 mg/kg) subcutaneously and MPO activity was determined using o-dianisidine method 48 h later. CN210 (20 and 50 mg/kg) and dexamethasone were administered 30 min before and 24 h after indomethacin treatment. Data are shown as mean ± SEM (n = 7). Statistically significant difference at \**P* < 0.05 from control (vehicle alone) and ^#^*P* < 0.05 from normal (without indomethacin) analyzed by one-way ANOVA with Holm-Sidak’s test.

**Fig 3.**
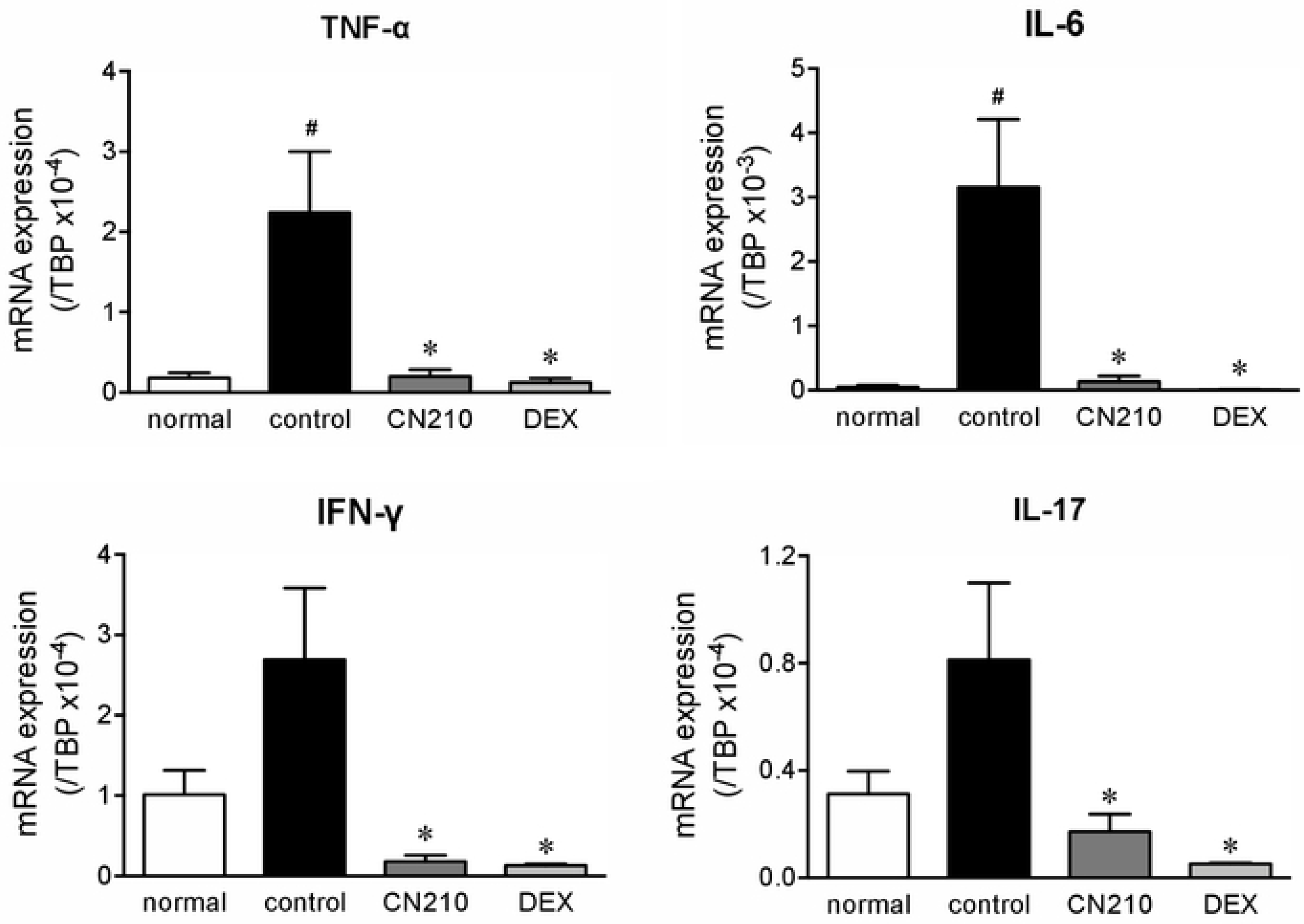
Effect of CN210 on indomethacin-induced increase in cytokine expression in the ileal mucosa. Animals received indomethacin (10 mg/kg) subcutaneously and mRNA expression for TNF-α, IL-6, IFN-γ, and IL-17 was determined by a quantitative real-time RT-PCR 48 h later. CN210 (20 and 50 mg/kg) and dexamethasone were administered 30 min before and 24 h after indomethacin treatment. Data are shown as mean ± SEM (n = 8). Statistically significant difference at \**P* < 0.05 from control (vehicle alone) and ^#^*P* < 0.05 from normal (without indomethacin) analyzed by one-way ANOVA with Holm-Sidak’s test.

### CN210 suppressed LPS-stimulated cytokine expressions in RAW264 cells

To further characterize the effects of CN210 on inflammatory cellular responses, we next examined the effects of CN210 in a monocyte/macrophage cell line, RAW264. The exposure of RAW264 cells to LPS (1 µg/mL) markedly upregulated the gene expression of cytokines such as TNF-α, IL-6, IL-12A, and IL-23A (Fig 4). These responses were suppressed by co-treatment with CN210 (0.01~10 μM), in a concentration-dependent manner, whilst significant effects taking place at a concentration as low as 0.01 µM.

**Fig 4.**
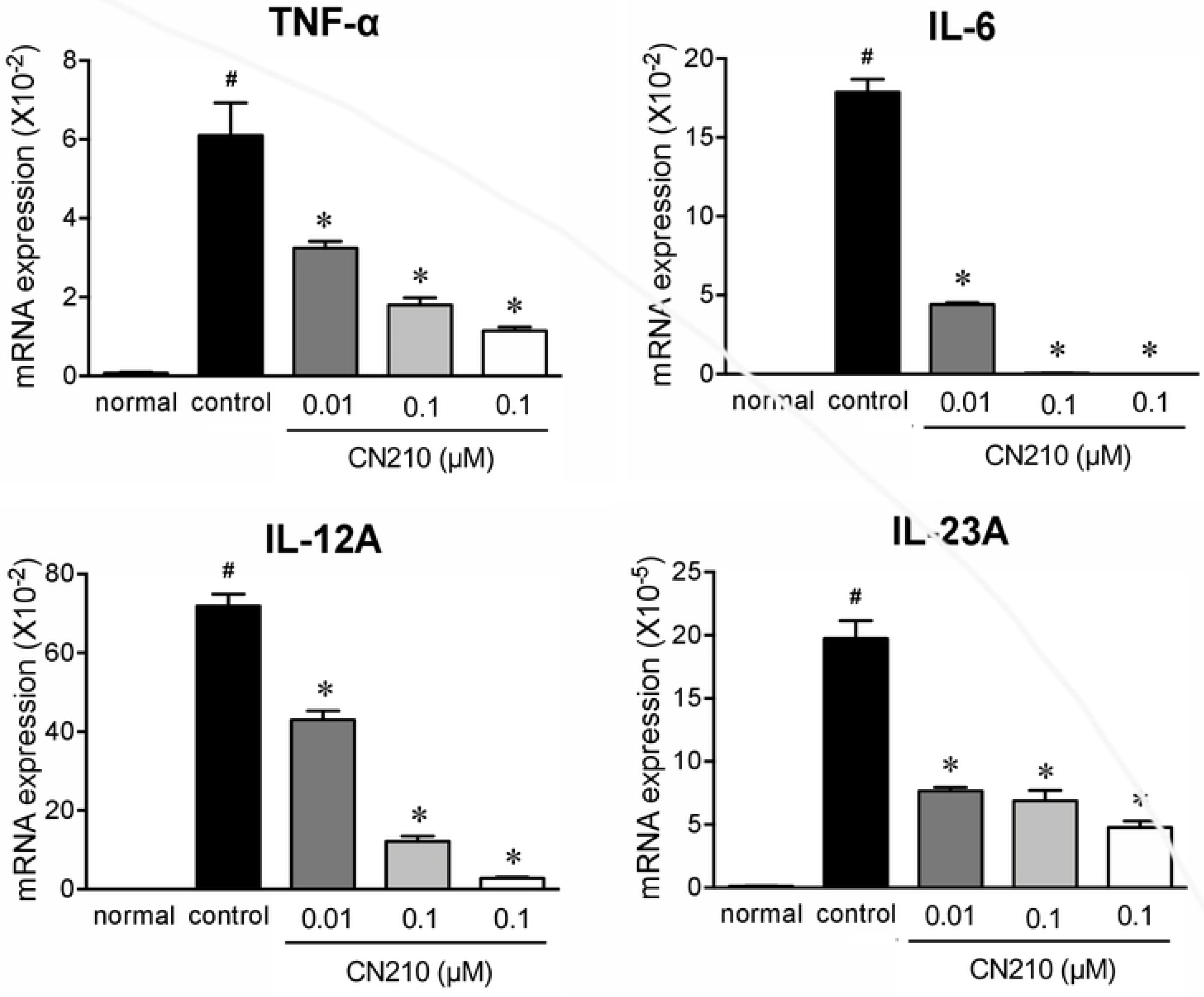
Effect of CN210 on LPS-stimulated cytokine expression in RAW264 cells. The cells were treated with LPS (1 µM) and total RNA was isolated 4 h later. CN210 (0.1–10 µM) were co-treated with LPS. indomethacin-induced increase in cytokine expression in the ileal mucosa. The mRNA expression of TNF-α, IL-6, IL-12A, and IL-23A was determined by a quantitative real-time RT-PCR 48 h later. Data are shown as mean ± SEM (n = 6). Statistically significant difference at \**P* < 0.05 from control (vehicle alone) and ^#^*P* < 0.05 from normal (without indomethacin) analyzed by one-way ANOVA with Holm-Sidak’s test.

### CN210 suppressed LPS-activated NF-κB, ERK, and JNK in RAW264 cells

To elucidate the involvement of intracellular signalling pathways in CN210-induced anti-inflammatory effects, we examined the effects of CN210 on NF-κB, ERK, p38 and JNK activations in RAW264 cells. The exposure of RAW264 cells to LPS (1 µg/mL) induced translocation of NF-κB p65 subunit to nuclei (Figs 5A and B). This translocation was impaired significantly by co-treatment with CN210 (10 µM). Likewise, LPS treatment enhanced phosphorylation of p38, ERK, and JNK in RAW264 cells. Whilst co-treatment with CN210 (10 µM) significantly suppressed phosphorylation of ERK and JNK, the suppressive effect was more prominent with ERK phosphorylation compared to JNK phosphorylation. Interestingly, CN210 did not suppress LPS-induced phosphorylation of p38 in RAW264 cells.

**Fig 5.**
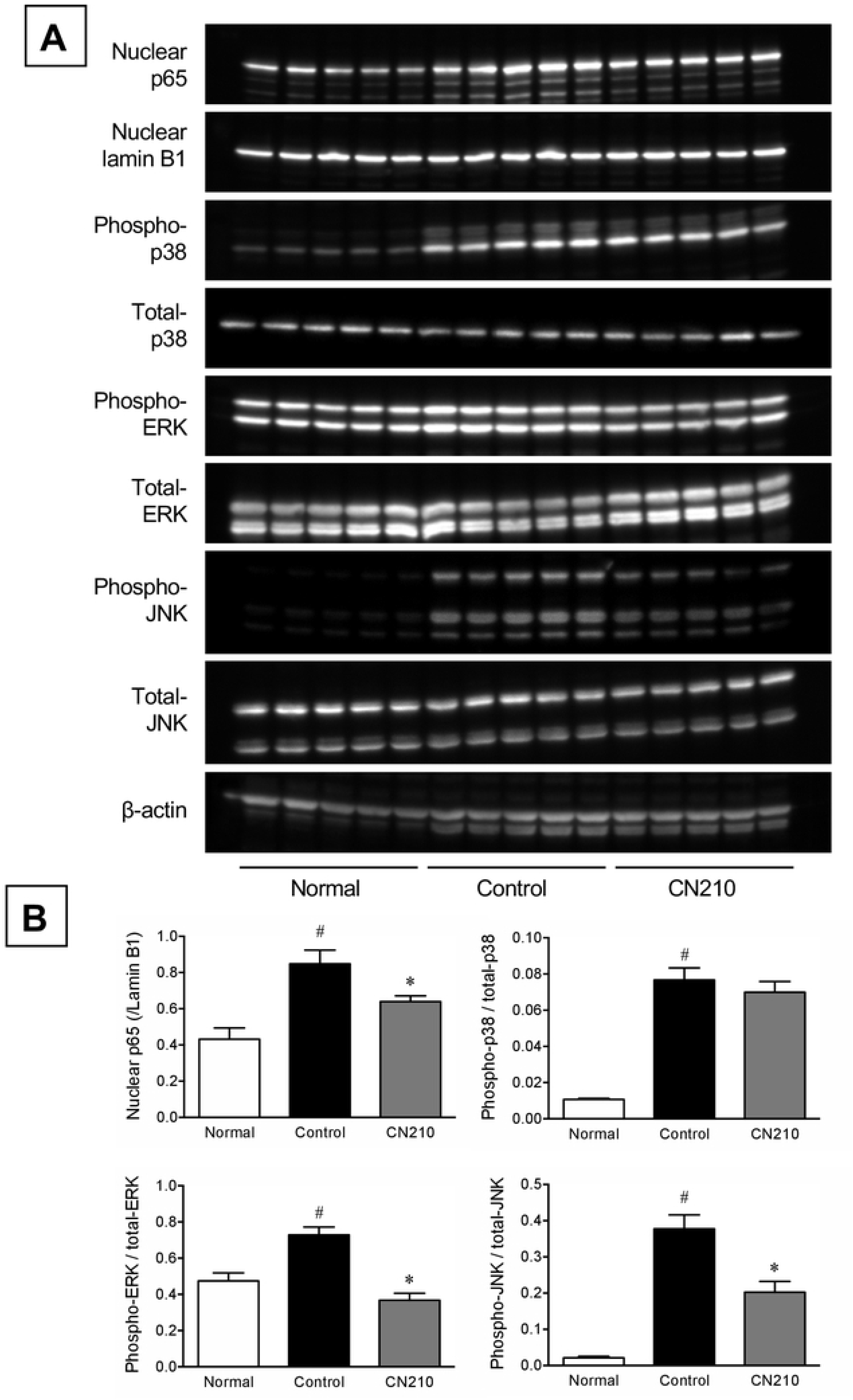
Effect of CN210 on LPS-activated NF-κB and MAPK in RAW264 cells. The cells were treated with LPS (1 µM) and activation of NF-κB and MAPK was determined by Western blotting 4 h later. CN210 (0.1–10 µM) were co-treated with LPS. animals. Data are shown as mean ± SEM (n = 5). Statistically significant difference at \**P* < 0.05 from control (vehicle alone) and ^#^*P* < 0.05 from normal (without indomethacin) analyzed by one-way ANOVA with Holm-Sidak’s test.

## Discussion

In recent years, BET inhibitors have attracted attention as a novel class of drug candidates not only for cancer but also for various inflammatory diseases [13, 14]. Several studies demonstrated that BET inhibitors were effective in the models of arthritis, vascular inflammation, acute pancreatitis, hepatic fibrosis and renal damage [23, 29–32], as well as in T cell-transfer colitis models [21, 22]. In the present study, we demonstrated that a novel quinazoline-based bromodomain inhibitor, CN210, inhibited CBP and p300 in addition to BET proteins and that CN210 potently attenuated indomethacin-induced CD-like ileitis in mice.

BET proteins, such as BRD2 and BRD4, epigenetically regulate transcription of pro-growth genes and pro-inflammatory cytokine genes [33]. Several reports showed that small-molecule BET inhibitors and siRNA suppressed cytokine production in inflammatory cells such as macrophages and synovial fibroblasts [29, 34]. Furthermore, it is known that BET proteins regulate gene transcriptions during the differentiation of naïve CD4^+^ cells to mature helper-T cells, and that BET inhibitors, such as JQ1, attenuated the differentiation of Th1, Th2, and Th17 cells [35, 36]. BET inhibitors have also reported to suppress inflammatory cytokine production and T-cell activation *via* inhibition of dendritic cell maturation and promotion of regulatory T-cell differentiation [22].

In the present study, we utilized an experimentally-induced murine CD-like ileitis model to assess CN210 as a potential drug candidate of CD. We demonstrated that CN210 interfered with the upregulation of the genes that encode TNF-α, IL-6, IFN-γ, and IL-17, which are known to play key roles in the pathogenesis of indomethacin-induced ileitis in in experimental animals [37–39]. Importantly, suppression of IFN-γ and IL-17 may indicate that CN210 may provide longer-term benefits by interfering T cell differentiation and dendritic cell maturation.

To study the anti-inflammatory effects of CN210 in more detail, we examined its effects on LPS-stimulated cytokine expression in monocyte/macrophage cell line RAW264 *in vitro*. Previous study showed that BET inhibitor, JQ1, and knockdown of BET family proteins with siRNA blocked cytokine productions in LPS-treated RAW264 cells [34]. CN210 potently suppressed the upregulation of the genes encoding TNF-α, IL-6, IL-12α, and IL-23α in RAW263 cells. Because IL-12 and IL-23 are important regulators for differentiation and maintenance of Th1 and Th17 cells, respectively [40, 41], CN210 may also provide prolonged therapeutic benefit by interfering with antigen-presented cell responses to induce differentiation and maintenance of Th1 and Th17 cells.

CN210 also reduced LPS-induced translocation of NF-κB p65 subunit (RelA) to nuclei (Figs 5A and B). It has been shown that acute stimulation of endothelial cells with TNF-α resulted in a rapid redistribution of chromatin activators guided by NF-κB forming *de novo* SE complex, which can be abrogated by JQ1 [42]. In addition, inhibition with JQ1 or knockdown of BRD2 and BRD4 genes with shRNA reduced NF-κB activation in response to TNF-α in human vascular endothelial cells [23]. Lastly, in addition to binding acetylated histones, BRD4 was shown to directly bind RelA acetylated by p300, thereby co-activating the genes involved in inflammatory response [43]. Taken together, the interfering effects of CN210 on NF-κB nuclear translocation may play a role in exerting its anti-inflammatory effects.

The MAPKs are known to comprize a family of serine/threonine kinases activated by diverse stimulations and induce various cellular functions including inflammatory responses. There are three major classes of MAPKs in mammals; ERK, p38, and JNK [44]. Previous studies demonstrated that BET inhibition reduced activation of all three types, *i.e.* p38, JNK, and ERK [23, 45]. In contrast, CN210 suppressed LPS-induced phosphorylation of JNK and ERK but not p38. The reason for this difference is unknown and warrants further investigation.

In the present study, we found that CN210 showed competitive binding effects for the bromodomains of CBP and p300 in addition to its activity against BET proteins (Table 1). CBP and p300 also are bromodomain-containing proteins and are paralogous histone acetyltransferases (HATs) and transcription co-activators [46, 47]. CBP and p300 play an important regulatory role in gene expression *via* NF-κB- and MAPK-dependent pathways, especially JNK-dependent pathways [48, 49]. In a study with spleen cells derived from mice that lack one allele of CBP, induction of TNF-α gene expression in T-cells in response to anti-CD3 antibody stimulation was shown to be impaired, implying a key role of CBP in T-cell immune response [50]. A recent study showed that SGC-CBP30, a CBP/p300-specific inhibitor, reduced immune cell production of IL-17A and other proinflammatory cytokines [51]. SGC-CBP30 also inhibited IL-17A secretion by Th17 cells from healthy donors and patients with ankylosing spondylitis and psoriatic arthritis [51]. Taken together, CBP and p300 are considered as emerging drug targets for inflammatory diseases and their inhibition afforded by CN210 may have contributed to potent anti-inflammatory effects of CN210. Additional studies, including the examination of the effects of CN210 on HAT activity of CBP and p300, as well as on the transcriptional machinery are ongoing.

## Conclusions

In conclusion, CN210, a bromodomain inhibitor of BET proteins and CBP/p300 proteins, attenuates indomethacin-induced CD-like ileitis in mice *via* inhibition of cytokine expression due at least in part to the suppression of NF-κB and MAPK activations and is a potential drug candidate for IBD with a unique mechanism of action.

## Author contributions

Conceived and designed the experiments: MY SK. Performed the experiments: TN KH SK SK. Analyzed the data: KM MY SK. Contributed reagents/materials/analysis tools: KM MY S-MY DJM SK. Wrote the paper: MY SK. Checked and revised the manuscript: S-MY.

## Supporting information

**S1 Raw images. Original blot images.** Full unedited images for Fig 5A.

